# Hydroxyl radical footprinting analysis of a human haptoglobin-hemoglobin complex

**DOI:** 10.1101/2021.09.20.461051

**Authors:** Dmitry S. Loginov, Jan Fiala, Peter Brechlin, Gary Kruppa, Petr Novak

**Affiliations:** BioCeV - Institute of Microbiology of the CAS, Prumyslova 595, CZ-252 50 Vestec, Czech Republic; Orekhovich Institute of Biomedical Chemistry, Pogodinskaja str. 10, 119191, Moscow, Russia; Faculty of Science, Charles University, Hlavova 8, CZ-128 20 Prague, Czech Republic; Bruker Daltonik GmbH, Fahrenheitstraße 4, 28359 Bremen, Germany; Bruker s.r.o., Prazakova 60, 619 00 Brno, Czech Republic

**Author notes:** To whom correspondence should be addressed (D. Loginov); (P. Novak).

**Keywords:** FPOP, footprinting, haptoglobin, hemoglobin, timsTOF Pro

## Abstract

Methods of structural mass spectrometry have become more popular to study protein structure and dynamics. Among them, fast photochemical oxidation of proteins (FPOP) has several advantages such as irreversibility of modifications and more facile determination of the site of modification with single residue resolution. In the present study, FPOP analysis was applied to study the hemoglobin (Hb) – haptoglobin (Hp) complex allowing identification of respective regions altered upon the complex formation. Oxidative modifications were precisely localized on specific residues using a timsTOF Pro mass spectrometer. The data allowed determination of amino acids directly involved in Hb – Hp interactions and those located outside of the interaction interface yet affected by the complex formation. Data are available via ProteomeXchange with identifier PXD021621.

## 1. Introduction

Currently, methods of structural mass spectrometry have become a powerful tool in the field of structural biology facilitating studies of protein folding and interactions. In this field, fast photochemical oxidation of proteins (FPOP) introduced in 2005 [1] provides valuable data related to a structural dynamics of proteins. This method is based on an irreversible covalent modification of amino acids side chains by hydroxyl radicals, which can be generated by a variety of methods including synchrotron radiolysis of water, electrochemistry, or laser photolysis of H_2_O_2_ [2]. The broad reactivity of hydroxyl radicals results in possible modifications of 19 amino acids, and 14 of these modifications are readily detected by mass spectrometry so that FPOP-MS analysis provides a rich and informative dataset related to chemical footprinting of the surfaces of proteins [3].

Reactivity of amino acids with ^•^OH varies by up to two orders of magnitude [3] meaning some difficulties may be encountered in detection of modified peptides with low relative abundance. Thus, developments in the instrumentation, e.g. increasing sensitivity of mass spectrometers, and improvements in peptide separation methods, etc., will allow more informative datasets to be obtained. From this point of view, a new mass spectrometer, the timsTOF Pro from Bruker Daltonics has features that should be very helpful. An integrated trapped ion mobility spectrometry (TIMS) adds a third dimension increasing the capacity for peptide separation [4], and the “Parallel Accumulation – Serial Fragmentation” (PASEF) method secures fast, reliable, and very sensitive fragmentation data from multiple precursors [5,6].

Hemoglobin (Hb) is one of the key proteins of the respiratory system, which transports oxygen and participates in the detoxification of reactive oxygen and nitrogen species [7]. Hb forms a tetramer by symmetric pairing of dimers consisting of the α- and β-globins, where each subunit contains a prosthetic heme molecule [8]. Erythrocyte-encapsulated Hb serves as a protector, but being released from them, it dissociates into αβ dimers and transforms into a very toxic protein due to the reactive properties of the heme [7]. To prevent possible toxic events an organism synthesizes an acute phase protein haptoglobin (Hp), which has the highest binding affinity for Hb [9]. In its turn, the Hb – Hp complex is recognized by the macrophage receptor CD163, which scavenges Hb [10]. The simplest Hp form is Hp 1-1 composed from two α- and two β-chains linked by disulphide bonds [11]. Currently, two X-ray structures of Hb – Hp complex are accessible in the PDB database. However, one structure represents complex of porcine proteins with a resolution 2.9 Å [12], and the second is a multimeric complex of human Hb – Hp with a receptor from *Trypanosoma brucei brucei* [13]. So far, there is not any high-resolution structural model of human Hb – Hp complex alone. Based on the structural data several amino acids participating in the Hb – Hp interactions have been indicated in porcine complex [12,14], yet the finest molecular details remains unclear for the human protein complex [15]. Thus, we applied FPOP analysis to study human Hb – Hp interactions in more details.

## 2. Materials and methods

Lyophilized human haptoglobin phenotype 1-1 (Hp) and human α,β haemoglobin (Hb) were purchased from Sigma Aldrich. Deglycosylation enzyme PNgase F was from New England BioLabs Inc. For digestion Typsin/Lys-C mass spectrometry grade proteases mixture from Promega Corporation was used. Additional chemicals reported in this article were purchased in the highest available purity from Merck.

### 2.1 Fast photochemical oxidation of proteins (FPOP)

Before the FPOP analysis the purity of the proteins was confirmed by SDS-PAGE electrophoresis (Supplementary fig. 1). Solutions of 0.27 mg/ml Hp, 0.27 mg/ml Hb, 7.5 mM H_2_O_2_ and 75 mM L-methionine were prepared in degassed 150 mM ammonium acetate (pH 6.8). The protein complex was formed by mixing Hp and Hb stock solutions in a volumetric ratio 1:1. The FPOP experiment was carried out in a continuous capillary flow system described in [16]. The system is composed of two syringe pumps (New Era, models NE-1000 and NE-4000) and three glass syringes (Hamilton) connected with quartz capillaries (Polymicro Technologies) and two MicroTees (Upchurch Scientific). Before the experiment, 250, 500 and 1000 μl syringes were filled with the protein sample, 7.5 mM H_2_O_2_ and 75 mM L-methionine, respectively. Flow rates were set to 10 ul/min, 20 ul/min and 20 ul/min, respectively. For the generation of hydroxyl radicals, a 248 nm KrF laser (Coherent, COMPex50) was used. The laser beam was focused on a quartz capillary with i.d. of 75 µm where a transparent window (approx. 6.5 mm) was formed by removal of the polyimide coating. Subsequently, the protein sample mixed with H_2_O_2_ was irradiated with one laser shot (15 Hz, 20 ns pulse duration, 2.24 mJ/mm^2^ radiant exposure) followed by quenching of hydroxy radicals with 75 mM L-methionine.

### 2.2 Protein digestion and LC-MS/MS analysis

Samples collected after the FPOP experiment were subjected to a reduction of cysteines with 20 mM Tris(2-carboxyethyl)phosphine, for 20 min, at 56 °C, followed by alkylation with 20 mM iodoacetamide for 20 min at 25 °C in the dark. Samples were then two times diluted with 100 mM 4-ethylmorpholine buffer (pH 8.5): acetonitrile (ACN) (90:10 v/v), followed by overnight deglycosylation with PNGase F (protein:enzyme ratio 1:20). Afterwards, trypsin/Lys-C was added (protein:enzyme ratio 1:20) and samples were digested for 8 hours at 37 °C. Digestion was stopped by addition of TFA to a final concentration of 0.1% and tryptic peptides were subsequently dried by SpeedVac (Eppendorf).

Samples were analyzed on an ultra-high pressure nanoflow chromatography system (nanoElute, Bruker Daltonics) coupled to a quadrupole time-of-flight mass spectrometer with a trapped ion mobility (TIMS) front end (timsTOF Pro, Bruker Daltonics) via a nano-electrospray ion source (Captive Spray Source, Bruker Daltonics). By microliter pickup, peptides (in amount corresponding to the 40 ng of digested proteins) were directly loaded and separated on an analytical column 25 cm x 75 µm, C18, 1.6 µm (Aurora Column, Ion Opticks, Australia). Peptides were eluted using 2 % ACN, 0.1% formic acid as mobile phase A at a flow rate of 400 nl/min, applying a 21 min gradient of 2-37% Acetonitrile by mobile phase B: ACN, 0.1% formic acid at 50 °C column oven temperature. The eluting peptides were interrogated by an MS acquisition method recording spectra from 100-1,700 m/z and ion mobility scanned from 0.6 to 1.6□Vs/cm^2^. The method consisted of a TIMS survey scan of 150 ms followed by 6 PASEF MS/MS scans, each 150 ms for ion accumulation and ramp time (ion accumulation is carried out in parallel, ions are accumulated for the next scan in the accumulation region of the TIMS device, while ions are being scanned out of the analyzer region). Total cycle time was 1.08 seconds. Target intensity was 40,000, intensity threshold 1000, singly charged peptides with m/z < 800 were excluded by an inclusion/exclusion polygon filter applied within the ion mobility – m/z heatmap. Precursors for data-dependent acquisition were fragmented with an ion mobility-dependent collision energy, which was linearly increased from 20 to 59□eV.

The raw data were deposited to the ProteomeXchange Consortium via the PRIDE partner repository with the dataset identifier PXD021621.

### 2.3 Data analysis

Acquired data were searched against a database containing the sequences of Hb and Hp (including deglycosylated form of β subunit) supplemented with sequences of common contaminants using PEAKS X+ software (Bioinformatics Solutions Inc., Waterloo, ON, Canada). Precursor ion tolerance was set at 10 ppm, and the mass tolerance for MS/MS fragment ions was set at 0.05 Da. Carbamidomethylation of cysteine and commonly observed FPOP modifications [3] were considered as variable modifications (Table 1).

**Table 1.** List of FPOP modifications.

| Modification | Targets |
| --- | --- |
| FPOP oxidation / + 15.994915 | M, W, Y, F, H, L, I, V, P, R, K |
| FPOP dioxydation / + 31.989828 | C, M, W, Y, F |
| FPOP trioxydation / + 47.984745 | C, W |
| FPOP carbonyl / + 13.979265 | L, I, V, P, R, E, Q, H |
| FPOP Met-32Da / - 32.008457 | M |
| FPOP His+5Da / 4.9789 | H |
| FPOP His-10Da / -10.031968 | H |
| FPOP His->Asp / -22.03197 | H |
| FPOP His->Asn / -23.01598 | H |
| FPOP Arg_deguanidination / - 43.053432 | R |
| FPOP decarboxylation / - 30.010565 | Q, N |
| FPOP ST_carbonyl / - 2.01565 | S, T |

Peptide-spectrum matches were filtered by peptide -10lgP scores ≥ 15, and threshold to a localization score assigned to modifications was set ≥ 20. FDR was set to 5 %. Intensities of peptides were determined using Compass DataAnalysis v.5.2 (Bruker Daltonics).

## 3. Results and discussion

Studying the molecular details of the human Hb – Hp interaction is a challenging task. Although both proteins have been subjected to various investigations and two 3D structures with resolution around 3 Å of their complex have been published [12,13], none of them represent a high resolution structural model of human Hb – Hp complex. Thus, methods of structural mass spectrometry might be useful in refining the proposed structural models. In the present study, FPOP was the method of choice, because of the deglycosylation step in the sample preparation, which is hardly compatible with HDX MS, an another common method used for protein structural studies [2]. Data from single proteins and their complexes were compared to determine regions and residues involved in the Hb – Hp interaction or altered by it.

Using a bottom-up approach a sequence coverage of Hpα and Hpβ was 100 % for the “free” protein and in the complex with Hb. Also, full sequence coverage was achieved for Hbα and Hbβ in the complex. For “free” Hb coverage was 59 % and 97 % for α and β subunits, respectively (Supplementary fig. 2-5).

Protein digests were measured using a recently introduced mass spectrometer, the timsTOF Pro with TIMS and PASEF technologies allowing an additional dimension of peptide separation (ion mobility) and fragmentation of a high number of precursors without loss of sensitivity [6]. Data were processed using PEAKS X+ software. A benefit of using PEAKS X+ is the possibility to introduce a high number of peptide modifications with reasonable computational time. Overall, 13 different modifications were included in the data search, including those occurring at multiple residues, like oxidation (+16 Da) of Phe, His, Ile, Lys, Leu, Met, Pro, Arg, Val, Trp, Tyr and +14 Da mass shift at Glu, His, Ile, Leu, Pro, Gln, Arg, and Val. Thus, we were able to get a very informative and rich dataset of identified peptides bearing up to three modifications (Table 2).

**Table 2.** Number of identified peptides by PEAKS X+ software.

| | Hbα | | | Hbβ | | | Hpα | | | Hpβ | | |
| --- | --- | --- | --- | --- | --- | --- | --- | --- | --- | --- | --- | --- |
|  | All | Mod. | Cert. | All | Mod. | Cert. | All | Mod. | Cert. | All | Mod. | Cert. |
| Hb | 149 | 78 | 34 | 218 | 142 | 50 | - | - | - | - | - | - |
| Hp | - | - | - | - | - | - | 89 | 29 | 2 | 282 | 116 | 58 |
| Hb-Hp | 294 | 144 | 49 | 335 | 184 | 52 | 169 | 59 | 2 | 429 | 184 | 61 |

All – all identified peptides; Mod. – number of modified peptides with a localization score ≥ 20; Cert. – number of modified peptides with at least one definite modified residue.

The high number of modified peptides arises from multiple possible modifications of the same residues. For example, six different modifications of His and Trp in mono-, di- and trioxidized states were found (Supplementary fig. 2-5). Taken together, these data allow precise footprinting of protein surfaces upon their interaction.

Quantitative analysis of modified peptides allowed the discovery of protein regions affected upon Hb – Hp complex formation (Fig. 1). Hpα was more oxidized (less protected) in the complex, including the C and N termini, while no significantly protected regions were observed. Rearrangements in the Hpβ structure upon the interaction with Hb resulted in protection of the N terminus and the region 271-311, and more exposure to oxidation of the C terminus and the region 203-270. On the contrary, protection of the C terminus and oxidation of the N terminus were observed in the case of Hbα. For Hbβ a more oxidized protein region upon complex formation was residues 105-146, including the N terminus. It should be mentioned that oxidation of the middle parts of Hbβ was detected only upon the complex formation (Fig 1c and 1d).

**Figure 1.**
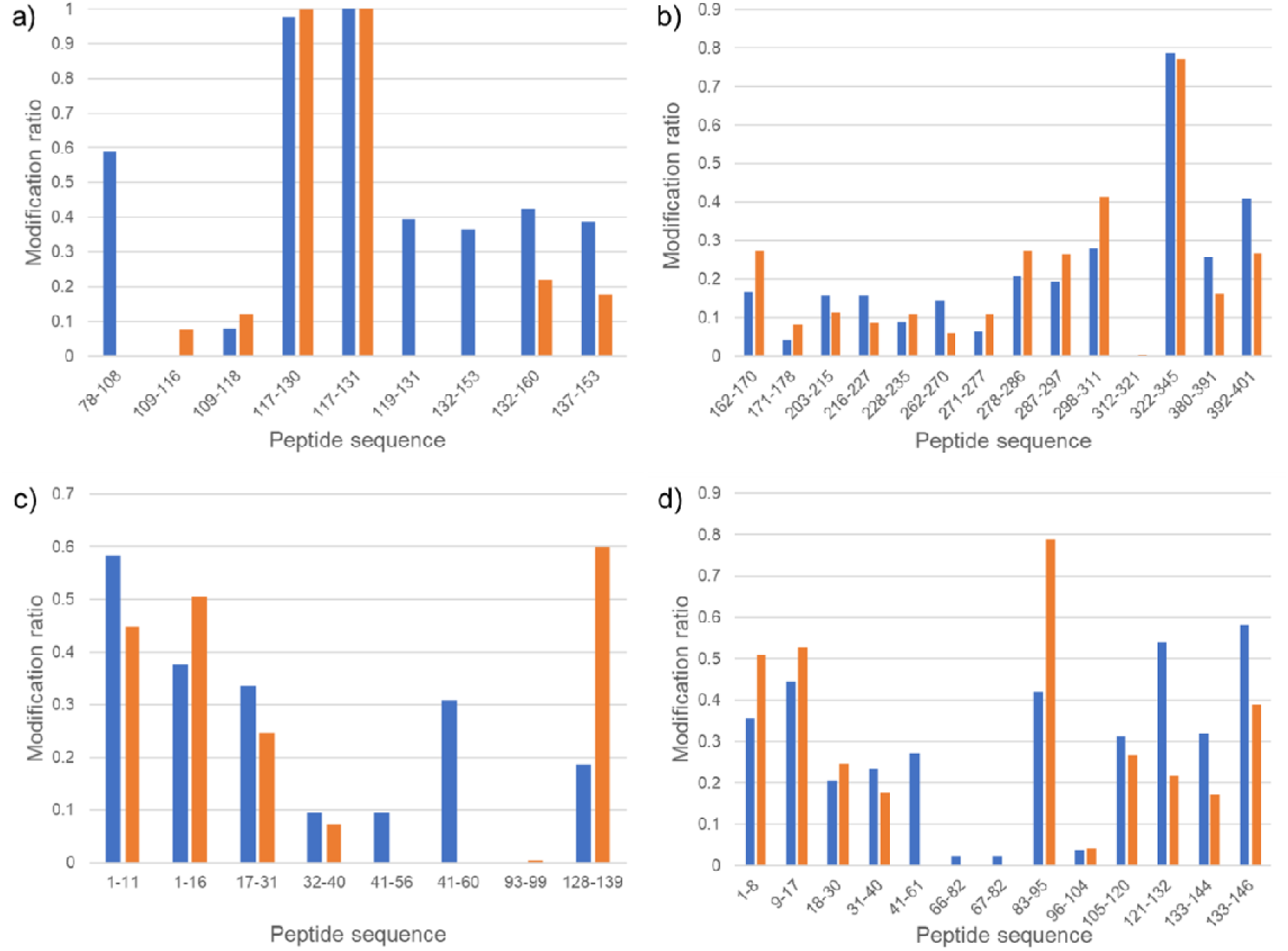
Extent of modification of peptides from Hb and Hp (Blue bars – Hb – Hp complex, orange bars – “free” protein). a – Hpα, b – Hpβ, c – Hbα, d – Hbβ.

One of the advantages of FPOP analysis over HDX is that single residue resolution is easier to achieve, as it is not necessary to resort to ETD or other non-ergodic fragmentation schemes to avoid scrambling of the labels. For this analysis we selected peptides with certainly determined modified residues and S/N ratio > 900 to obtain unambiguous data. Overall, an oxidation rate was determined for 2, 37, 27 and 32 residues in Hpα, Hpβ, Hbα and Hbβ, respectively (Fig. 2). Both certain modifications identified for Hpα corresponded to the trioxidation of cysteines. Among them, Cys111, the residue engaged in a disulfide bridge formation, that connects two αβHp dimers [12], was found to be susceptible to oxidation in the complex at a higher rate.

**Figure 2.**
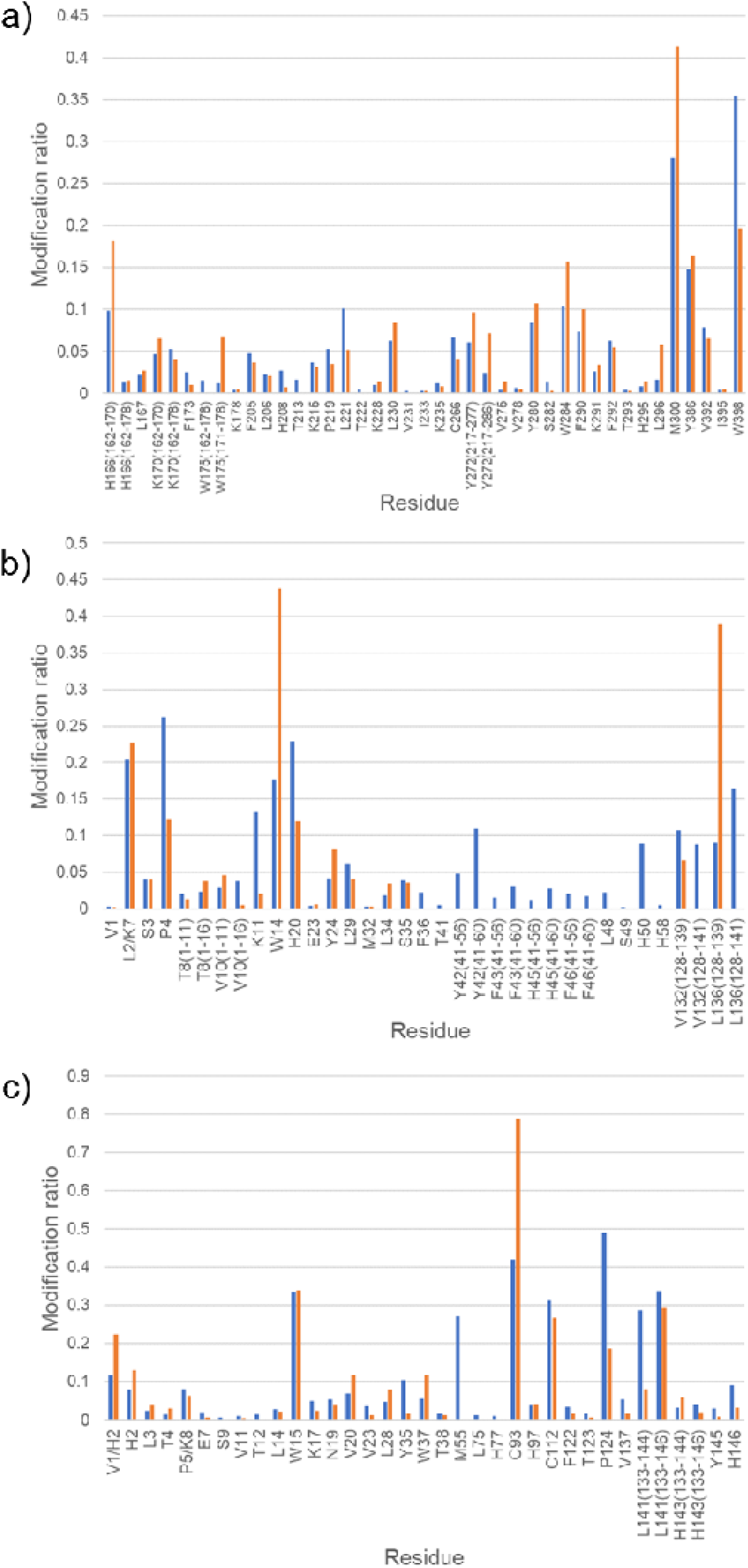
Modification ratios at the residue level (Blue bars – Hb – Hp complex, orange bars – “free” protein). a – Hpβ, b – Hbα, c – Hbβ. For the residues with multiple modifications, like His, the sum of all detected modifications contributing to the overall oxidation extent is presented (see Supplementary Table 1 for the respective values for the each modification).

Previously, several sites of Hb oxidation by H_2_O_2_ have been described [17,18]. The proposed hot spots included Tyr42 in the α subunit, Cys93 and Tyr145 in the β subunit. Also, His2, Trp15, Pro36, Trp37, Met55 and Cys112 in the Hbβ were found to be minor oxidative sites. Protection (decreased rate or absence of an oxidation) of all these sites upon Hb – Hp complex formation has also been shown [18]. Evidence for modification of all these residues but βPro36 were found during our FPOP analysis. Because of differences in the reaction conditions we observed modification of the neighboring residue βTyr35 which is a more favorable amino acid for the oxidation by ^•^OH [3], instead βPro36.

Evidence for strong protection of these Hb residues against oxidation upon interaction with Hp found in the previous study was not confirmed by our data. αTyr42 and βMet55 were found to be readily oxidized in the complex. None of the residues previously reported to be susceptible to oxidation by H_*2*_O_*2*_ were found to be completely protected upon Hb – Hp complex formation (Fig. 2). This discrepancy could be explained by the conditions of the radical generation. In Pimenova et al radicals were generated by Hb itself, thus no oxidation was found in the case of Hb – Hp complex [18]. In the present study radicals were generated by laser induced homolytic cleavage of H_2_O_2_.

Nine residues found to be modified during our FPOP analysis have been shown to be potentially involved in Hb – Hp interactions (Hb: αVal1, βTrp37, βHis97 and βTyr145 ; Hp: βLeu167, βLeu206, βPhe290, βLys291 and βHis295) [12]. No significant changes in the modification ratio of these residues in Hpβ were found (Fig. 2a). However, the overall oxidation level of the respective peptides, except 203-215, was lower in the complex, meaning the respective protein parts are protected due to Hb – Hp interactions (Fig. 1b). Residues 203-215 represents an unstructured region located outside of the interaction interface. Thus, its higher oxidation rate might be explained by some conformational changes in the complex favorable to the subsequent oxidation of amino acids side chains by hydroxyl radicals. Two Hb residues, namely αVal1 and βHis97, were oxidized at the same level as in the “free” protein (Fig. 2).

Previously βTrp37 and βPhe292 in Hb and Hp, respectively, have been proposed as contributing the most to the binding energy of Hb – Hp complex, so called hot spots, and interacting with each other based on the protein-protein docking calculation using structure of porcine Hb – Hp complex as the template [14]. The last was not confirmed by our data. The Hb βTrp37 residue was significantly protected upon the complex formation, whereas Hp βPhe292 was oxidized at the same level (Fig. 2). Thus, Hb βTrp37 is indeed involved in intermolecular interactions, but not with Hp βPhe292. This finding is in agreement with the X-ray structural model of human complex, where the Cα-Cα distance between these residues is 18.5Å making the interaction unlikely [13]. Further Phe292 is located on the surface of Hpβ in a near proximity of two complex glycan moieties (Asn184 and Asn207) protecting it from the possible complex formation [19].

Analysis of oxidation extents of different residues allowed the determination of the most affected residues upon the complex formation (Fig. 3). As expected, a majority of the protected residues were located in the Hb – Hp interaction interfaces, whereas the more oxidized ones were on the protein surfaces. Differences in the oxidation rates of Hp residues were less profound than in Hb. That is not surprising because Hb is subjected to more structural changes upon Hb – Hp complex formation. The Hb residues with increased oxidation rate located on the surface of α (Pro4, Lys11, Tyr42 and His50) and β (Met55 and Pro124) subunits might be considered as participating in the formation of Hb quaternary structure.

**Figure 3.**
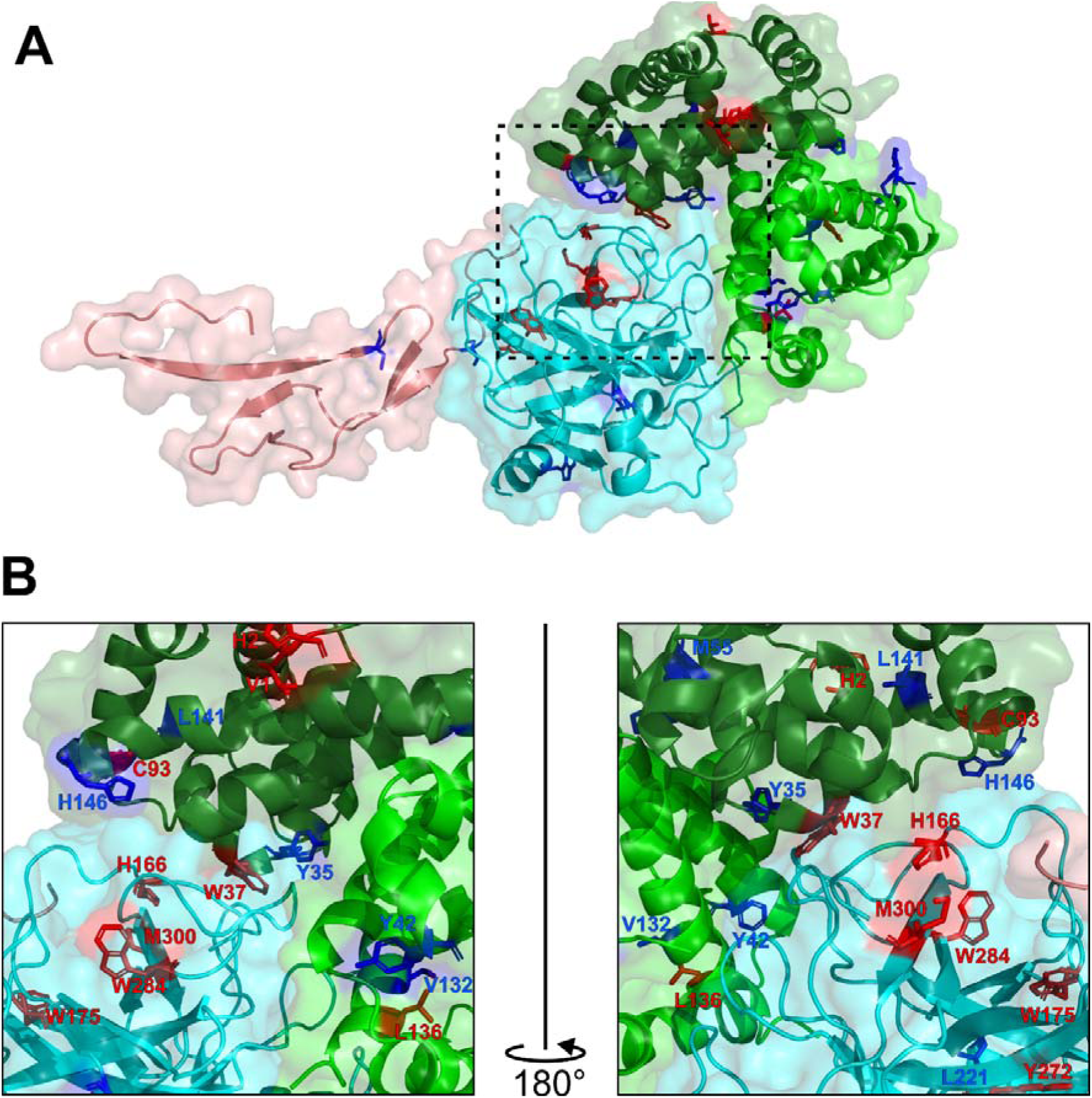
Structural model of Hb – Hp complex with highlighted residues oxidized by the FPOP experiment. a – Overview of a tetrameric complex composed of α,βHp and α,βHb subunits. b – Zoom of interaction interface. Red residues were found to be less oxidized (protected) upon the formation of complex, showing evidence of the interaction interface. Blue residues represent amino acids that were more oxidized after the complex’s formation. Hpα is in pink, Hpβ is in cyan, Hbα is in green and Hbβ is in dark green. Respective light colors represent solvent accessible area of the protein surface.

## 4. Conclusions

FPOP analysis enables an efficient structural analysis of Hb – Hp interactions. The advantage of using FPOP for studying of protein-protein interactions is the possibility to follow even minor conformational changes providing a complete dataset related to a possible interaction mechanism. The FPOP data allowed determination of regions in the protein structures affected upon the complex formation, and single residues involved in the interactions were identified. Three Hpβ residues located in the interaction interface, namely His166, Trp284 and Met300, were shown to be protected upon the complex formation. Also, lower oxidation rates were determined for αLeu137, βTrp37 and βCys94 in Hb. These data indicate involvement of the residues in the Hb – Hp interaction. The FPOP dataset also identifies Hb residues participating in the formation of its quaternary structure.

## Supporting Information

Supplementary figure 1. SDS-PAGE electrophoresis of human α,β hemoglobin and human haptoglobin phenotype 1-1. Lane 1 – hemoglobin, lane 2 – haptoglobin, Std – protein ladders.

Supplementary figure 2. Peptide map of haptoglobin α. A – HpHb complex, B – free protein. Modifications localized on the particular residue from fragmentation spectra are indicated above the sequence line.

Supplementary figure 3. Peptide map of haptoglobin β. A – HpHb complex, B – free protein. Modifications localized on the particular residue from fragmentation spectra are indicated above the sequence line.

Supplementary figure 4. Peptide map of hemoglobin α. A – HpHb complex, B – free protein. Modifications localized on the particular residue from fragmentation spectra are indicated above the sequence line.

Supplementary figure 5. Peptide map of hemoglobin β. A – HpHb complex, B – free protein. Modifications localized on the particular residue from fragmentation spectra are indicated above the sequence line.

Supplementary table 1. List of multiple modifications detected on the same residues.

## Acknowledgements

We thank Dr. Filip Dycka (University of South Bohemia, Czech Republic) for providing access to the PEAKS software and we acknowledge the Centre of molecular structure Core Facility at BIOCEV, a facility funded by European Regional Development Funds (CZ.1.05/1.1.00/02.0109 BIOCEV) and sup-ported by the Czech Infrastructure for Integrative Structural Biology (Structural mass spectrometry CF - LM2018127 CIISB for CMS BIOCEV funded by MEYS CR).

## References

[1] T.T. Aye, T.Y. Low, S.K. Sze, Nanosecond laser-induced photochemical oxidation method for protein surface mapping with mass spectrometry, Anal. Chem. 77 (2005) 5814–5822. https://doi.org/10.1021/ac050353m.

[2] X.R. Liu, M.M. Zhang, M.L. Gross, Mass Spectrometry-Based Protein Footprinting for Higher-Order Structure Analysis: Fundamentals and Applications, Chem. Rev. (2020). https://doi.org/10.1021/acs.chemrev.9b00815.

[3] G. Xu, M.R. Chance, Hydroxyl radical-mediated modification of proteins as probes for structural proteomics, Chem. Rev. 107 (2007) 3514–3543. https://doi.org/10.1021/cr0682047.

[4] J.A. Silveira, M.E. Ridgeway, M.A. Park, High resolution trapped ion mobility spectrometery of peptides, Anal. Chem. 86 (2014) 5624–5627. https://doi.org/10.1021/ac501261h.

[5] F. Meier, S. Beck, N. Grassl, M. Lubeck, M.A. Park, O. Raether, M. Mann, Parallel accumulation-serial fragmentation (PASEF): Multiplying sequencing speed and sensitivity by synchronized scans in a trapped ion mobility device, J. Proteome Res. 14 (2015) 5378–5387. https://doi.org/10.1021/acs.jproteome.5b00932.

[6] F. Meier, A.D. Brunner, S. Koch, H. Koch, M. Lubeck, M. Krause, N. Goedecke, J. Decker, T. Kosinski, M.A. Park, N. Bache, O. Hoerning, J. Cox, O. Räther, M. Mann, Online parallel accumulation–serial fragmentation (PASEF) with a novel trapped ion mobility mass spectrometer, Mol. Cell. Proteomics. 17 (2018) 2534–2545. https://doi.org/10.1074/mcp.TIR118.000900.

[7] I.K. Quaye, Extracellular hemoglobin: The case of a friend turned foe, Front. Physiol. 6 (2015) 1–13. https://doi.org/10.3389/fphys.2015.00096.

[8] A.N. Schechter, K. Singer, H. Lehmann, W. Castle, T. Huisman, E. Jaffe, ASH 50th anniversary review Hemoglobin research and the origins of molecular medicine, Blood. 112 (2008) 3927–3938. https://doi.org/10.1182/blood-BLOOD.

[9] P. Ascenzi, A. Bocedi, P. Visca, F. Altruda, E. Tolosano, T. Beringhelli, M. Fasano, Hemoglobin and heme scavenging, IUBMB Life. 57 (2005) 749–759. https://doi.org/10.1080/15216540500380871.

[10] M. Kristiansen, J.H. Graversen, C. Jacobsen, O. Sonne, H.J. Hoffman, S.K.A. Law, S.K. Moestrup, Identification of the haemoglobin scavenger receptor, Nature. 409 (2001) 198–201. https://doi.org/10.1038/35051594.

[11] F. Polticelli, A. Bocedi, G. Minervini, P. Ascenzi, Human haptoglobin structure and function - A molecular modelling study, FEBS J. 275 (2008) 5648–5656. https://doi.org/10.1111/j.1742-4658.2008.06690.x.

[12] C.B.F. Andersen, M. Torvund-Jensen, M.J. Nielsen, C.L.P. De Oliveira, H.P. Hersleth, N.H. Andersen, J.S. Pedersen, G.R. Andersen, S.K. Moestrup, Structure of the haptoglobin-haemoglobin complex, Nature. 489 (2012) 456–459. https://doi.org/10.1038/nature11369.

[13] K. Stødkilde, M. Torvund-Jensen, S.K. Moestrup, C.B.F. Andersen, Structural basis for trypanosomal haem acquisition and susceptibility to the host innate immune system, Nat. Commun. 5 (2014) 1–8. https://doi.org/10.1038/ncomms6487.

[14] C. Nantasenamat, V. Prachayasittikul, L. Bulow, Molecular Modeling of the Human Hemoglobin-Haptoglobin Complex Sheds Light on the Protective Mechanisms of Haptoglobin, PLoS One. 8 (2013). https://doi.org/10.1371/journal.pone.0062996.

[15] S. Tamara, V. Franc, A.J.R. Heck, A wealth of genotype-specific proteoforms fine-tunes hemoglobin scavenging by haptoglobin, Proc. Natl. Acad. Sci. U. S. A. (2020). https://doi.org/10.1073/pnas.2002483117.

[16] X.R. Liu, D.L. Rempel, M.L. Gross, Protein higher-order-structure determination by fast photochemical oxidation of proteins and mass spectrometry analysis, Nat. Protoc. 15 (2020) 3942–3970. https://doi.org/10.1038/s41596-020-0396-3.

[17] Y. Jia, P.W. Buehler, R.A. Boykins, R.M. Venable, A.I. Alayash, Structural basis of peroxide-mediated changes in human hemoglobin: A novel oxidative pathway, J. Biol. Chem. 282 (2007) 4894–4907. https://doi.org/10.1074/jbc.M609955200.

[18] T. Pimenova, C.P. Pereira, P. Gehrig, P.W. Buehler, D.J. Schaer, R. Zenobi, Quantitative mass spectrometry defines an oxidative hotspot in hemoglobin that is specifically protected by haptoglobin, J. Proteome Res. 9 (2010) 4061–4070. https://doi.org/10.1021/pr100252e.

[19] P. Pompach, K.B. Chandler, R. Lan, N. Edwards, R. Goldman, Semi-automated identification of N-glycopeptides by hydrophilic interaction chromatography, nano-reverse-phase LC-MS/MS, and glycan database search, J. Proteome Res. 11 (2012) 1728–1740. https://doi.org/10.1021/pr201183w.

