## Supplementary figures and images for "Hydroxyl radical footprinting analysis of a human haptoglobin-hemoglobin complex"

### Supplemental figure 1

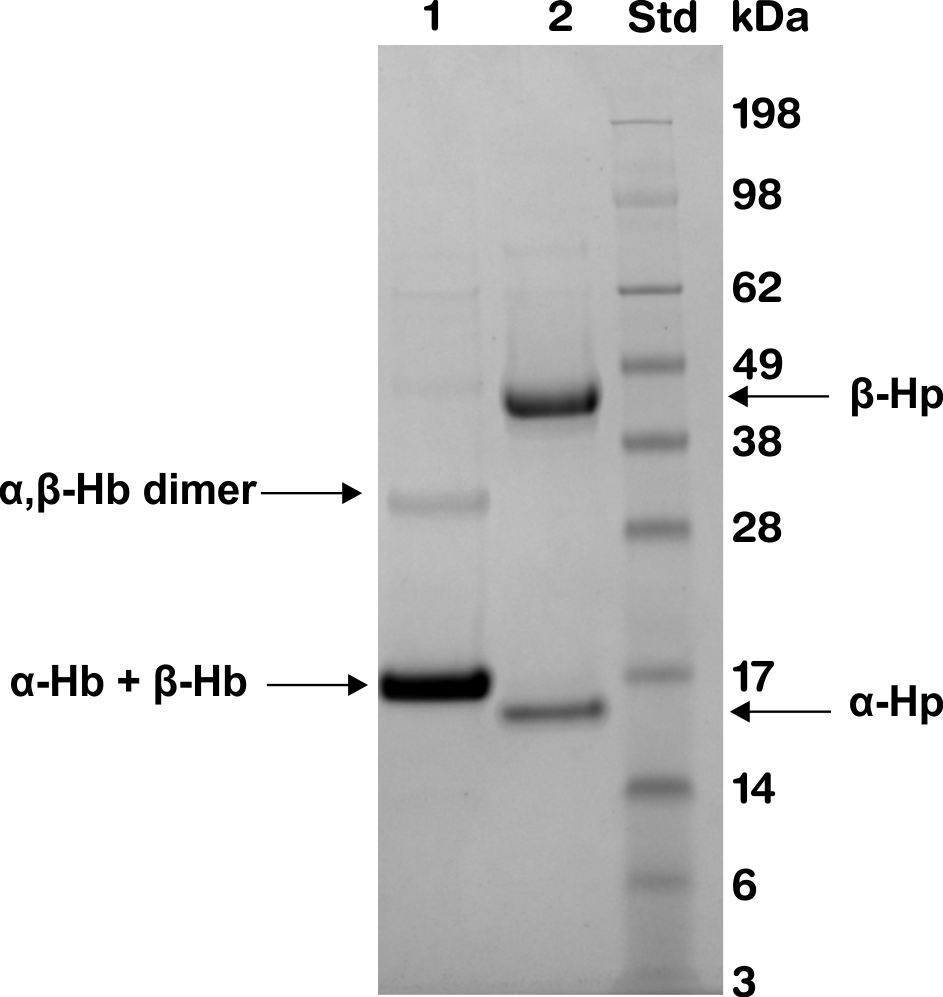

### Supplemental figure 2

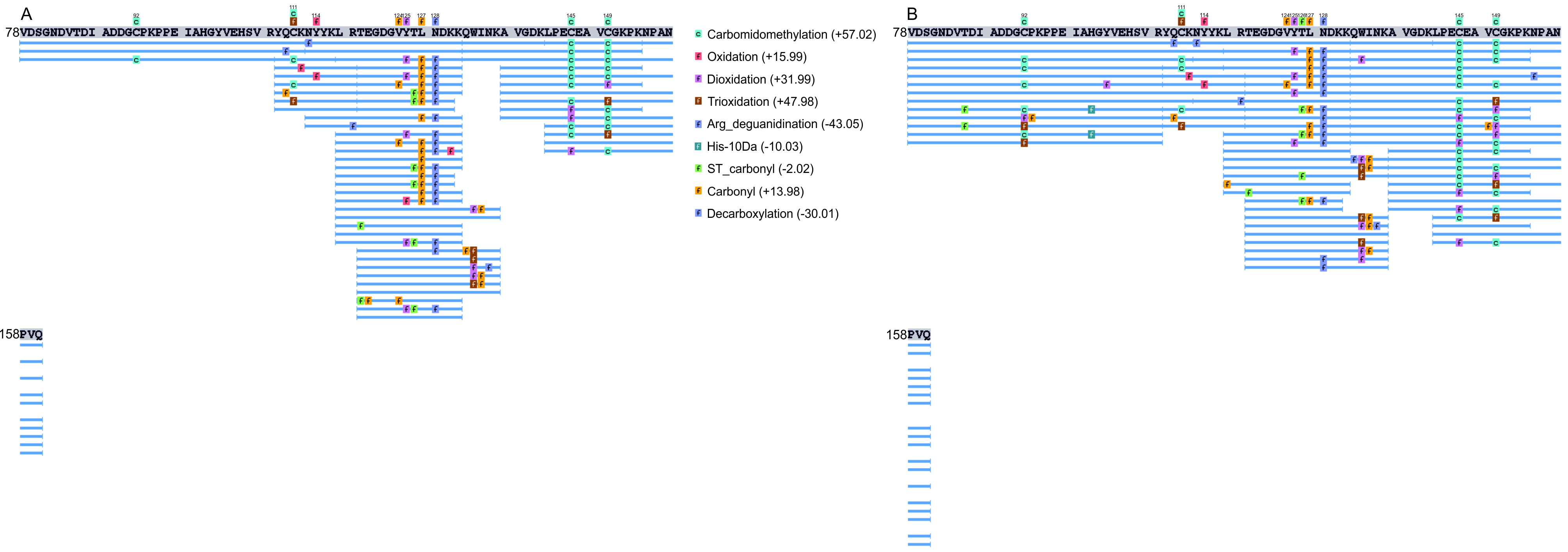

### Supplemental figure 3

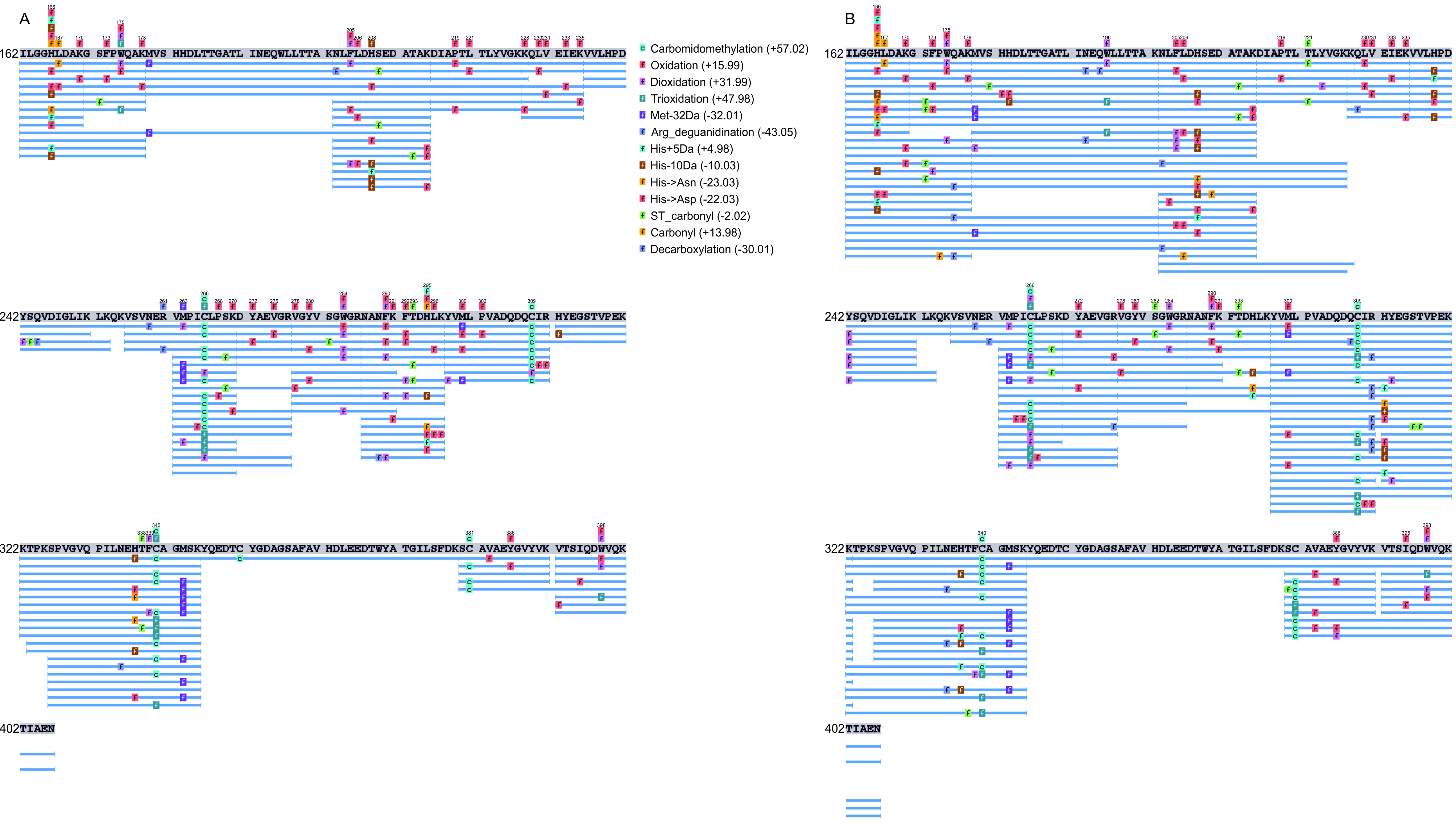

### Supplemental figure 4

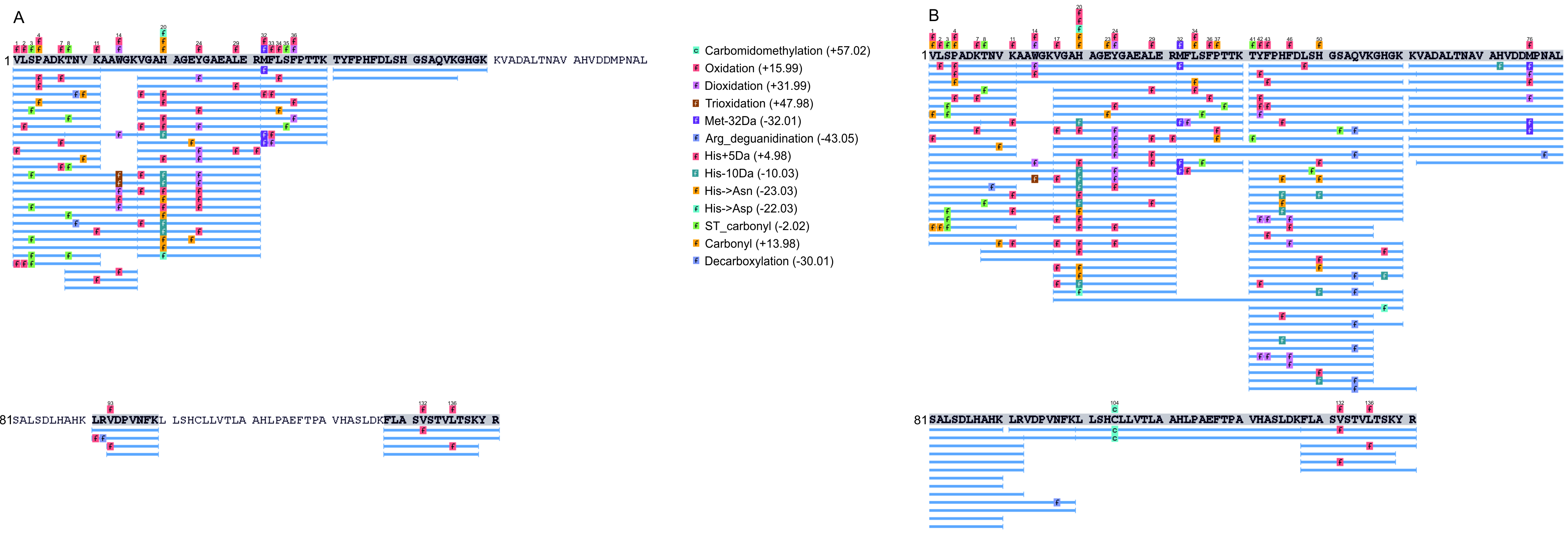

### Supplemental figure 5

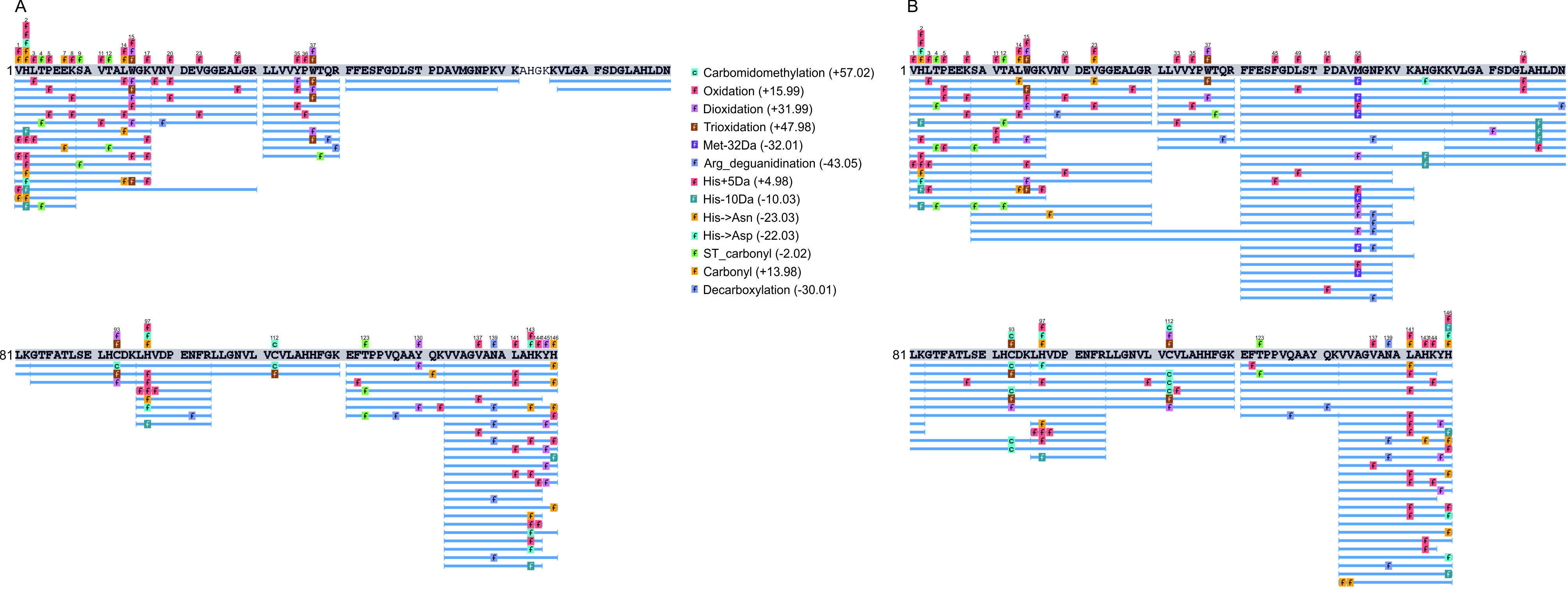
